# Flow Cytometric Evaluation of Yeast-Bacterial Cell-Cell Interactions

**DOI:** 10.1101/2021.10.13.464218

**Authors:** Ming Lei, Vikas D. Trivedi, Nikhil U. Nair, Kyongbum Lee, James A. Van Deventer

## Abstract

Synthetic cell-cell interaction systems can be useful for understanding multicellular communities or for screening binding molecules. We adapt a previously characterized set of synthetic cognate nanobody-antigen pairs to a yeast-bacteria coincubation format and use flow cytometry to evaluate cell-cell interactions mediated by binding between surface-displayed molecules. We further use fluorescence-activated cell sorting (FACS) to enrich for a specific yeast-displayed nanobody within a mixed yeast-display population. Finally, we demonstrate that this system supports characterization of a therapeutically relevant nanobody-antigen interaction: a previously discovered nanobody that binds to the intimin protein expressed on the surface of enterohemorrhagic *E. coli*. Overall, our findings indicate that the yeast-bacteria format supports efficient evaluation of ligand-target interactions. With further development, this format may facilitate systematic characterization and high throughput discovery of bacterial surface-binding molecules.

## Introduction

Surface components of bacterial cells such as cell membrane or cell wall proteins are an emerging class of therapeutic targets.^*1*^ For example, pathogenic bacteria present virulence factors on their cell surfaces in response to an external physiological stimulus to promote adherence to gut epithelial cells.^*2*^ Similarly, in the context of biofilm formation, various cell surface adhesion and colonization factors can participate in adhesion by selectively binding to extracellular matrix proteins on host cells such as fibronectin, collagen, and mucin.^*3*^ Strategies to target surface components responsible for virulence or adhesion include using 1) antibodies against polysaccharides and adhesins; and 2) antimicrobial peptides and antibiotics in conjunction with extracellular polymeric substance (EPS) matrix inhibitors.^*4*^ Traditional methods for discovery of surface component-binding antibodies and peptides have employed screens utilizing soluble, purified forms of a target protein of interest with cells displaying a library of binding molecules^*5*^; or conversely, a soluble library of binding molecules with a surface-anchored target protein.^*6*^ In addition to these well-established methods, recent studies have shown that cell-cell interaction assays can facilitate the discovery of novel binding molecules that recognize targets presented on cell surfaces. To date, these efforts have focused on utilizing microbial display platforms to discover binding proteins that recognize specific components of mammalian cell surfaces. Salema et al. directly screened an immune nanobody library against the epidermal growth factor receptor (EGFR), a protein overexpressed on the surface of human tumor cells, in an *E. coli* display format using magnetic cell sorting (MACS) and were able to isolate high affinity EGFR binders.^*7*^ In another notable example, Yang et al. used a yeast-mammalian cell-cell interaction system coupled with fluorescence-activated cell sorting (FACS) to discover antibodies against a human mu opioid G protein-coupled receptor by incubating a *S. cerevisiae* yeast display library with mammalian cells that were overexpressing the receptor. The yeast and mammalian cells expressed intracellular green fluorescent protein (EGFP) and red fluorescent protein (mCherry) reporters, respectively, enabling FACS of yeast-mammalian cell complexes. The study showed that formation of yeast-mammalian cell complexes was significantly enhanced by the interaction of surface displayed molecules.^*8*^ Multiple groups have quantitatively analyzed yeast-mammalian cell interaction systems with adherent and suspension cultured mammalian cells, including investigations into the roles of binding affinity, surface density of yeast-displayed ligands, surface density of mammalian-displayed ligands, selection conditions, and linker lengths.^*9-12*^ Recent studies by Rao and coworkers have demonstrated that a yeast-yeast interaction format can also enable enrichment of cognate binder-target interactions^*13*^, discovery of protein-protein interactions, and quantitative estimation of binding affinities.^*14*^

In addition to their use in ligand discovery, cell-cell interaction systems support the generation and characterization of synthetic interactions mediated by surface-displayed proteins or cell surface modifications. Glass et al. designed a set of *E. coli* cell-cell interactions modulated by orthogonal interactions between surface displayed nanobody-antigen pairs and characterized the formation and patterning of cell aggregates using microscopy and cell culture optical density.^*15*^ Similarly, a proof-of-concept synthetic *E. coli* cell-cell interaction system engineered using the highly stable SpyTag/SpyCatcher interaction pair was found to generate cell aggregates capable of activating a quorum sensing circuit.^*16*^ In a synthetic model system of bacterial interaction with human cells, modifications to the cell surface of *E. coli* using oligodeoxynucleotide-small molecule conjugates enabled engineered cells to physically interact with human cells overexpressing a cognate receptor.^*17*^ Finally, Chun et al. showed that yeast displaying a phage endolysin exhibited antimicrobial activity against *Staphylococcus aureus*,^*18*^ with bacterial lysis being an indirect indication of yeast-bacteria interactions since the endolysin was tethered to the yeast surface. Further adaptation of these synthetic interaction systems to yeast display format has the potential to enable new screening methodologies for discovering novel binding or antibacterial molecules that may be generalizable towards different bacterial species. A yeast-based discovery platform could have significant advantages over other formats, such as the lack of susceptibility of yeast to many antimicrobial peptides. As an initial step towards developing this platform, it would be useful to evaluate a model system of cell-cell interactions between yeast and *E. coli*. We are not aware of any previous studies performing this type of characterization.

In this study, we use flow cytometry assays to evaluate yeast-bacteria cell-cell interactions mediated by surface-displayed nanobody-antigen pairs. Initial investigations use interaction pairs previously implemented in an *E. coli*-*E. coli* interaction system described by Glass et al.^*15*^ Following display of nanobodies on the *S. cerevisiae* surface and antigens on the *E. coli* surface, we use flow cytometry to determine that coincubation of yeast and *E. coli* displaying cognate nanobody-antigen pairs results in significantly increased interaction events. These yeast-bacteria interaction events can further be used to enrich for a cognate, yeast-displayed nanobody from a mixed yeast population incubated with *E. coli* displaying the target antigen using fluorescence-activated cell sorting (FACS). The flow cytometry assay format is also able to detect binding events mediated by the interaction between a previously discovered nanobody and intimin, a virulence factor and potential therapeutic target expressed on the surface of enterohemorrhagic *E. coli* (EHEC).^*6, 19*^ The detection of this therapeutically relevant interaction in yeast-*E. coli* format indicates that this format facilitates identification of physiologically relevant interactions. Overall, the characterizations of yeast-bacteria cell-cell interactions performed in this study demonstrate the utility of this system for evaluating ligand-target interactions on cell surfaces. These results have the potential to support future development of a new screening platform for discovery of bacterial surface-binding molecules.

## Results and Discussion

### Yeast-bacteria cell complex formation is driven by specific interactions mediated by surface-displayed molecules

To investigate the potential for yeast and *E. coli* to form complexes driven by specific interactions, we used a previously described set of nanobody (Nb)-antigen (Ag) pairs known to exhibit mutual specificity to one another. In prior work, both the nanobodies and the antigens were displayed on the surfaces of *E. coli*.^*15*^ Here, we moved the nanobodies into a standard yeast (*S. cerevisiae*) display format (see *Materials and Methods* for details). Nanobodies are covalently anchored on the yeast surface using the Aga1p-Aga2p system, in which a heterologous protein fused to the α-agglutin subunit Aga2p is covalently anchored to the cell wall protein Aga1p.^*20*^ Yeast displaying Nb1, Nb2, or Nb3 were stained with calcofluor white (CFW), a blue fluorescent dye that stains the chitin in yeast cell walls and minimally stains gram-negative bacteria such as *E. coli*. Antigens were anchored on the *E. coli* surface using the intimin N-terminus from EHEC O157:H7.^*15*^ *E. coli* displaying Ag1, Ag2, or Ag3 were stained with SYTO 9, a green fluorescent dye that permeates cell membranes and binds to nucleic acids. Since SYTO 9 can stain both yeast and *E. coli* cells, SYTO 9-stained *E. coli* cells were washed with phosphate buffered saline (PBS) before coincubation with yeast to remove excess unbound dye and minimize the detection of false positive events. Though the specificity, half-life, and toxicity of such fluorescent dyes should be considered, a major advantage of using fluorescent dyes is that they can facilitate flow cytometry analysis without the need to genetically engineer cells to express fluorescent reporter proteins. This can expand the range of cell types that can be tested in this cell-cell interaction format to include genetically intractable organisms.

For reproducible evaluation of cell-cell interaction events, we experimentally determined several important assay parameters, including cell densities, cell ratios, and incubation time. We determined that a total cell density of 0.5 to 1 (OD_600_) gave reproducible results with a low level of nonspecific binding events in flow cytometry analysis. A reliable yeast:*E. coli* cell ratio for quantifying cell complex formation and minimizing nonspecific binding events in this system was approximately 1:1 (Figure S1). We observed that nonspecific interactions between yeast and *E. coli* increased as the relative amount of *E. coli* increased. The green fluorescence corresponding to SYTO 9 staining was stable for up to two hours for the conditions used in this study’s cell-cell aggregation assays.

We next evaluated the extent to which the two cell types, when co-incubated, associated with one another as a function of different interaction pairs (Figure 2A). Figure 2B depicts two-dimensional flow cytometry dot plots used to evaluate yeast and *E. coli* cell complexes. High levels of blue and green fluorescence are expected when flow cytometry events include at least one yeast cell and at least one *E. coli* cell. Upon coincubation of yeast and *E. coli* displaying Nb2 and Ag2, respectively, the frequency of events exhibiting high levels of both blue and green fluorescence increased (∼30-40% of total events) relative to coincubation of cells displaying noncognate nanobody-antigen pairs (∼10-15% of total events). A similar trend was observed with yeast displaying Nb3 and *E. coli* displaying Ag3. These results suggest that cognate Ab-Nb pairs mediate specific cell-cell interactions, resulting in detection of events on the flow cytometer that possess high levels of both blue and green fluorescence. We did not observe any increases in double-positive fluorescent events upon coincubation of Nb1-displaying yeast and Ag1-displaying *E. coli*. A contributing factor may be that, in our hands, *E. coli* cells induced to express Ag1 tended to form aggregates with other Ag1 expressing *E. coli* cells. Further, the OD600 of these cultures was less than *E. coli* cultures induced to express Ag2 or Ag3. This tendency to aggregate may interfere with potential yeast-*E. coli* interactions. In any case, our data strongly suggest that the formation of yeast-*E. coli* complexes can be mediated by specific sets of interactions.

**Figure 1.**
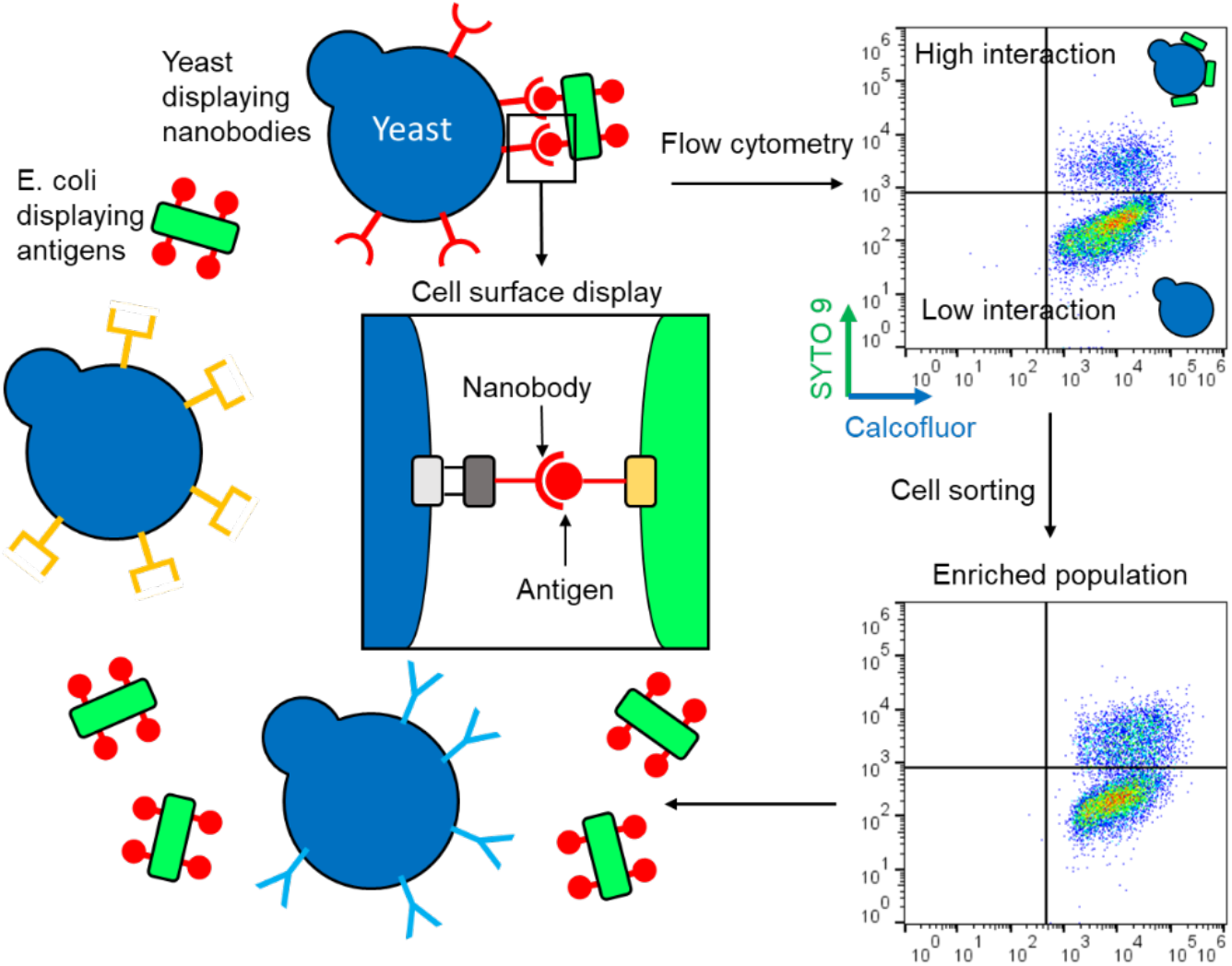
Overview of yeast-bacteria interaction system. Molecules displayed on the surface of yeast and *E. coli* can interact with each other through cognate binding. Flow cytometric analysis facilitates evaluation of interactions, and fluorescence-activated cell sorting enables enrichment for yeast displaying a specific binding partner.

**Figure 2.**
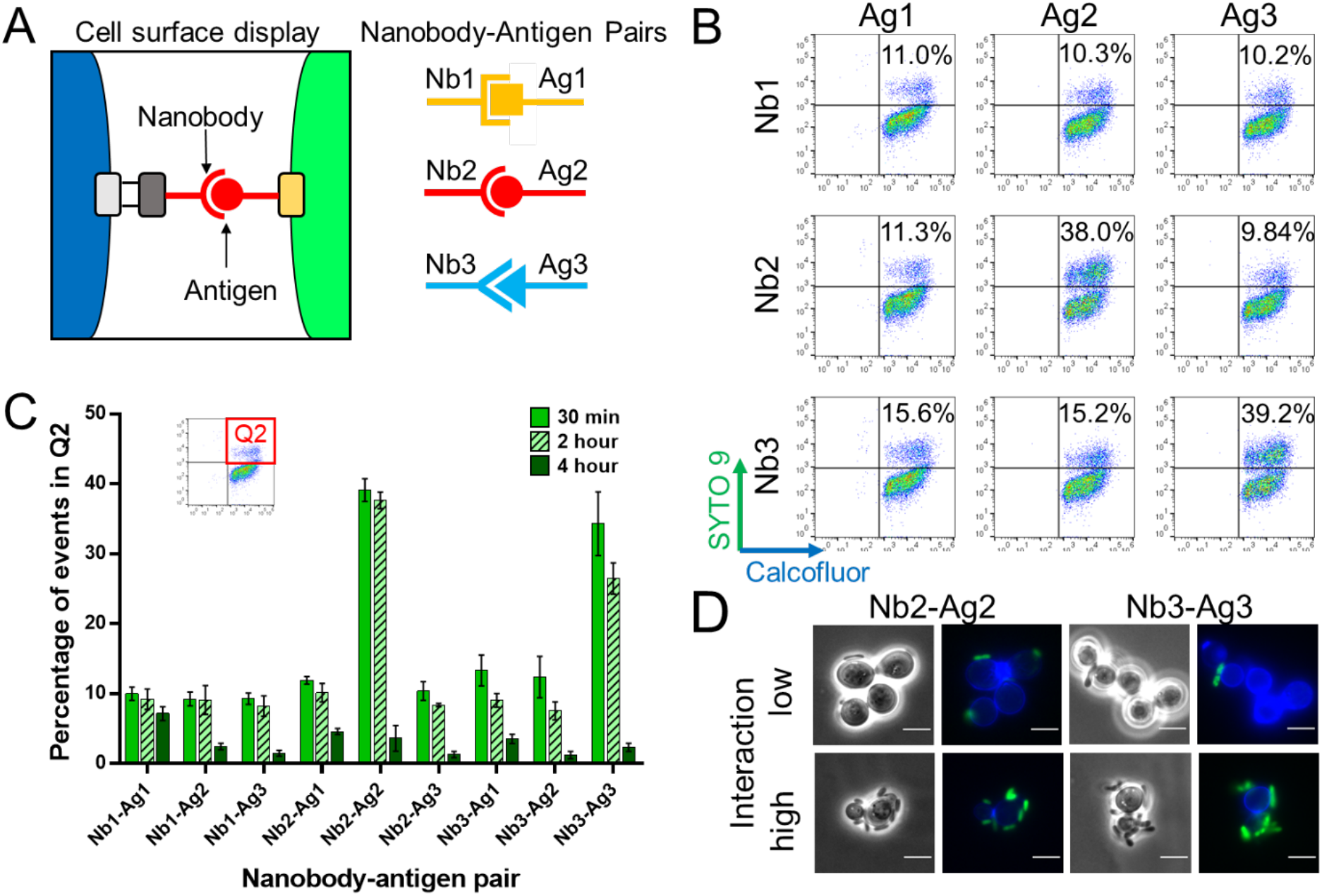
Flow cytometry and microscopy characterization of yeast-bacteria cell complexes. A) Schematic of orthogonal interactions between nanobodies and antigens displayed on yeast and *E. coli*, respectively. B) Representative flow cytometry dot plots for all combinations of coincubations for yeast cells displaying Nb1, Nb2, or Nb3 with *E. coli* cells displaying Ag1, Ag2, or Ag3. C) Percentage of events in quadrant 2 (Q2, high levels of blue and green fluorescence detected) following coincubation for 30 min, 2 hours, and 4 hours. Error bars represent the standard deviation of 3 biological replicates. D) Representative phase contrast and overlaid fluorescence microscopy images of low and high interactions between calcofluor-stained yeast displaying Nb2 or Nb3 (blue) and SYTO 9-stained *E. coli* displaying Ag2 or Ag3 (green) cells within the same samples. Scale bars indicate 5 µm.

To visualize yeast-*E. coli* complexes for evidence of direct physical interactions between the two cell types, cell suspensions were collected after a 30-minute coincubation and analyzed using fluorescence microscopy. A total OD of 2 was used for this analysis due to technical limitations; the density of cells previously used for flow cytometry was too sparse for capturing representative images of cells on the microscope within a single field of view. Yeast-*E. coli* complexes were observed for all interaction pairs, including non-cognate interactions. While some yeast cells induced to express the cognate nanobody had multiple *E. coli* cells bound, others had few or none, suggesting that the yeast cells exhibited varying display densities. These observations were consistent with flow cytometry analysis, which measured nonspecific interactions in 10-15% of the total events for non-cognate nanobody-antigen pairs. Additionally, some yeast cells did not display nanobodies on the cell surface after induction, as evidenced by a lack of HA and c-Myc labeling for full length nanobody display in a portion of each induced yeast population (Figure S2). Microscopy images for cell complexes representative of both low and high interaction events are shown for the Nb2-Ag2 and Nb3-Ag3 interaction pairs (Figure 2D). It was not possible to quantitatively correlate the microscopy data to the flow cytometry measurements due to limitations on the number of images that could be taken within the lifetimes of the fluorescent dyes. The SYTO 9 dye was useful for quantifying cell-cell interactions up to 2 hours, but could not be reliably used for this purpose for longer times due to gradual loss of fluorescence. While microscopy can provide evidence of physical interactions between yeast and *E. coli*, it is less suited for quantitative analysis, further motivating the need for a flow cytometry-based assay to characterize cell-cell interactions.

### A cognate nanobody-antigen interaction can be enriched through FACS

After developing a flow cytometry assay for characterization of yeast-bacteria cell-cell interactions, we attempted to enrich for a cognate binding molecule in this interaction format. Previous studies have shown that this type of enrichment is possible using bacteria-mammalian cell,^*7*^ yeast-mammalian cell,^*8*^ yeast-yeast,^*13*^ and phage-mammalian cell screens.^*21*^ However, to the best of our knowledge, enrichment of cognate binding has not been demonstrated for a high-throughput screen in a yeast-bacteria cell-cell interaction format. In a proof-of-concept experiment, we sorted for nanobody-antigen interaction events from a coincubation culture of yeast and *E. coli*. Yeast populations, prepared at 1:1:1, 10:1:10, and 100:1:100 ratios of cells displaying Nb1, Nb2, and Nb3, respectively, were screened against *E. coli* displaying Ag2. Cell-cell interactions were quantified on an analytical flow cytometer prior to cell sorting (Figure 3A). The samples were then sorted by enriching for events exhibiting high levels of green fluorescence (Figure S3; We selected green fluorescence because the available cell sorter lacks a suitable laser for the excitation and detection of calcofluor white). Sorted populations were recovered under conditions that inhibited bacterial growth (yeast growth media supplemented with penicillin/streptomycin). No bacteria were observed in microscopic examination of the recovered cultures. After the enriched cultures were grown to saturation, they were diluted and induced for nanobody display. Cell-cell interactions between the enriched yeast populations and Ag2-displaying *E. coli* were quantified again on the analytical flow cytometer (Figure 3A). Although the pre-sort and post-sort samples were evaluated on different days, we observed that the percentages of binding events for single nanobody display and mixed ratio controls were consistent across independent experiments (Figure S4A), allowing for direct comparisons between pre-sort and post-sort populations. The increased percentages of double-positive fluorescent events observed via flow cytometry suggested that there was substantial enrichment of yeast displaying Nb2 from 1:1:1 and 10:1:10 mixed populations of Nb1:Nb2:Nb3 (Figure 3B). However, detectable enrichment of yeast displaying Nb2 did not occur from the 100:1:100 mixture. The percentages of double-positive fluorescent events in control yeast-bacteria populations containing individual yeast clones displaying Nb1, Nb2, or Nb3 are depicted alongside the pre-sort and post-sort samples (Figure 3B). The percentage of Nb2-displaying yeast in the population was estimated as follows: Nb1-, Nb2-, and Nb3-displaying yeast were mixed at predetermined ratios, incubated with Ag2-displaying *E. coli*, and the percentage of interaction events was measured to generate a calibration curve for the expected fraction of Nb2-displaying yeast versus percentage interaction events (Figure S4B). To corroborate the results of the flow cytometry analysis, we performed amplicon sequencing on yeast plasmid DNA isolated from pre-sort and post-sort samples to quantify the percentage of each of the three nanobody genes detected in each sample. The sequencing results indicated enrichment of Nb2 clones from all three starting ratios (Figure 3C). Additionally, there was a significant (p-value = 0.0004) linear correlation between the enrichment values calculated using flow cytometry analysis and sequencing (Figure 3D). On the other hand, the two assays yielded different fold changes in enrichment. This difference can be attributed to technical limitations of both assays. The flow cytometry assay measures the percentage of double positive interaction events for a given population of co-incubated yeast and *E. coli* cells, which includes nonspecific cell-cell interactions. The DNA sequencing estimates the number of genes present in the full population, including cells in the population that did not exhibit detectable protein display at the time of flow cytometry analysis (Figure S2). Nevertheless, the cell-cell interaction system enabled enrichment of a yeast-displayed nanobody to its cognate antigen presented on the *E. coli* surface; this was confirmed using both flow cytometry and sequencing analysis.

**Figure 3.**
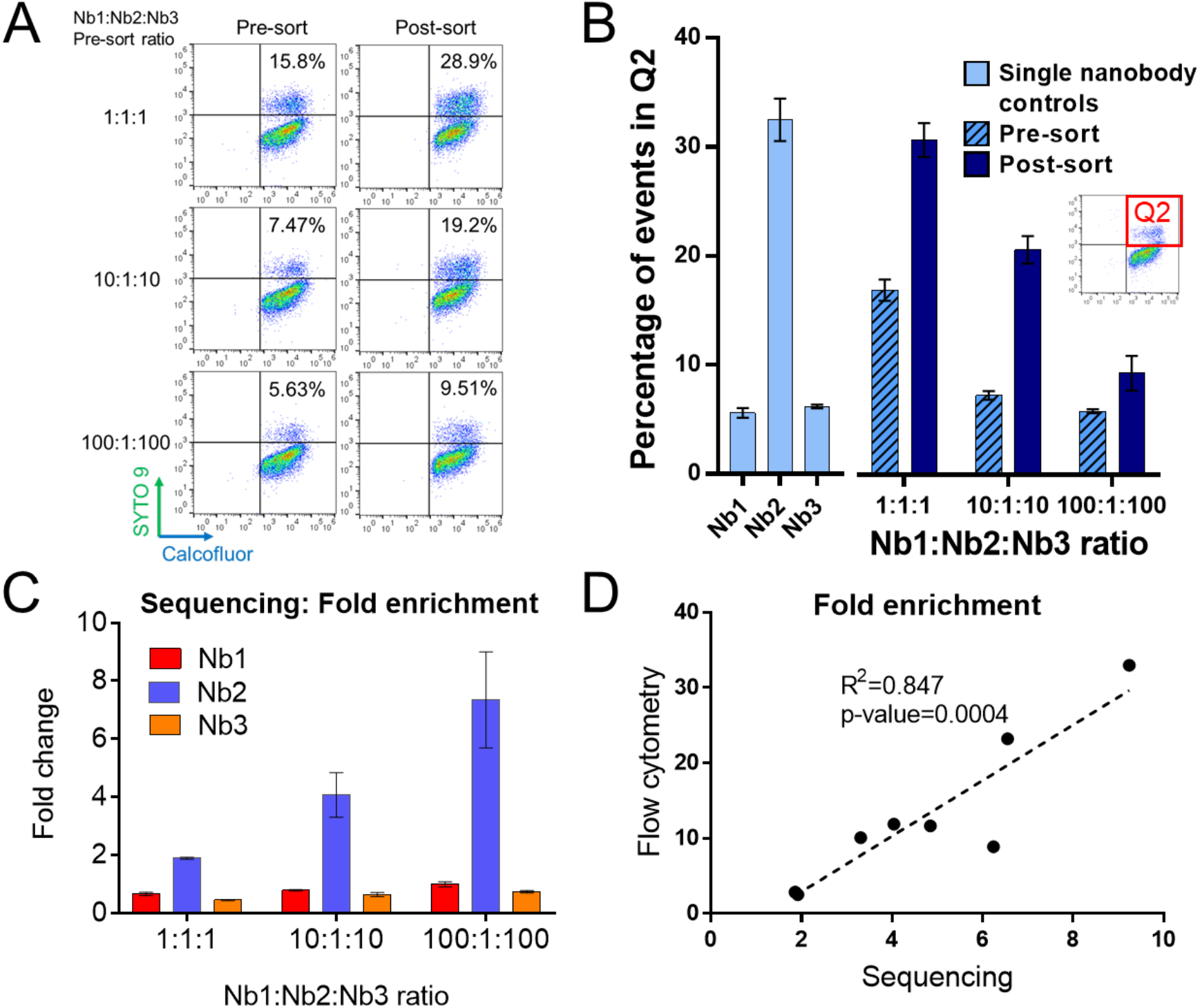
Cell sorting on mixed yeast display-nanobody populations against an *E. coli*-displayed target antigen. A) Dot plots of coincubations containing pre- and post-sort yeast populations. B) Percentage of events in Q2 for incubation of Ag2-expressing *E. coli* with control yeast clones each displaying a single nanobody and with yeast populations comprised of mixed ratios of cells displaying one of three nanobodies, pre- and post-sort. Error bars represent the standard deviation of 3 technical replicates sorted and processed independently for downstream steps. C) Fold change of sequencing reads in pre-sort and post-sort samples. Error bars represent the standard deviation of 3 technical replicates for cell sorting. D) Correlation of fold enrichment values determined by flow cytometry analysis versus amplicon sequencing.

### Yeast-bacteria assay can detect a therapeutically relevant nanobody-antigen interaction

Intimin is a protein antigen expressed on the surface of enterohemorrhagic *E. coli* (EHEC) that mediates bacterial attachment to host tissues through interaction with the translocated intimin receptor, Tir. A previously characterized nanobody against intimin, IB10, was demonstrated to reduce intimin interaction with Tir.^*19*^ We evaluated whether surface display of the IB10-intimin nanobody-antigen pair would result in detectable enhancements in cell-cell interactions in our system.

We observed a significant increase in the frequency of events exhibiting high levels of both blue and green fluorescence for the cell-cell interaction events mediated by the IB10-intimin nanobody-antigen interaction pair relative to the Ag2 and DH5α controls (Figure 4), suggesting that this model system can be applied towards biologically relevant antigen targets. We also tested the ability of IB10-displaying yeast to bind directly to EHEC O157:H7, a strain that Is known to express intimin on its cell surface. Despite using conditions reported to upregulate expression of the EHEC virulence gene cluster that includes the intimin gene,^*22*^ we did not detect significant binding of yeast displaying the IB10 nanobody to EHEC as measured by our cell-cell interaction assay (data not shown). However, this observation is not surprising given that expression of EHEC pathogenicity genes is regulated by a complex genetic circuit that responds to environmental cues and integrates signals from host cells.^*22*^ In this regard, intimin overexpression on the *E. coli* surface can still serve as a biologically relevant model for intimin induction during host cell infection.^*23*^

**Figure 4.**
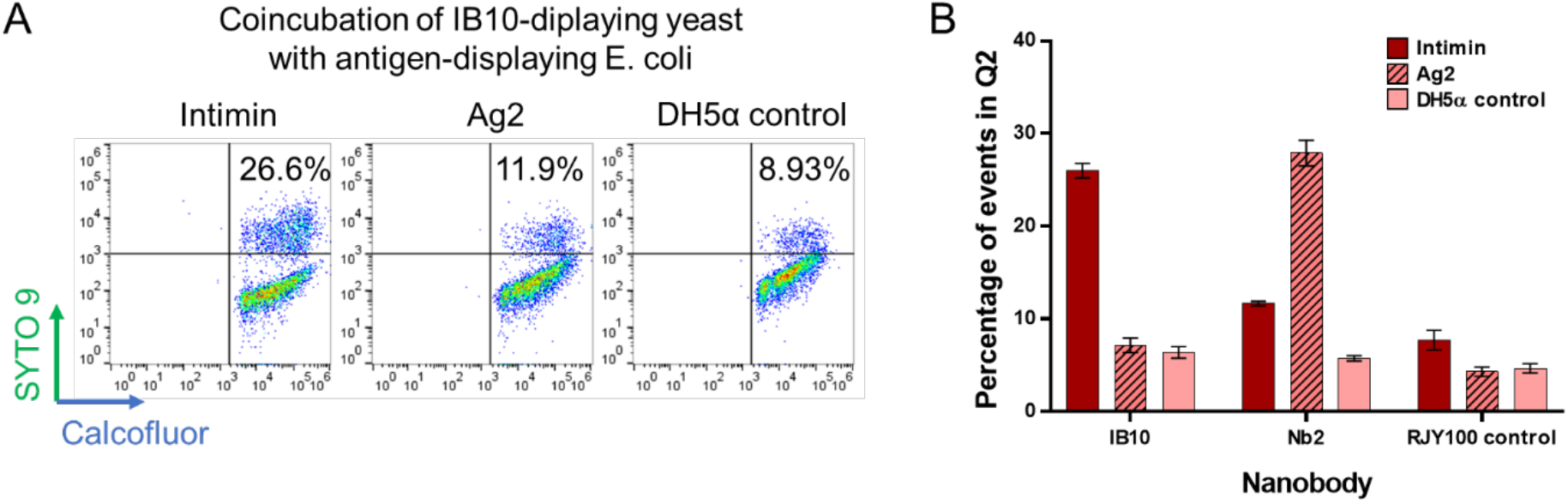
Application of interaction model system for surface-displayed intimin and anti-intimin nanobody. A) Representative dot plots for yeast-*E. coli* coincubations for yeast displaying IB10 with *E. coli* displaying intimin or Ag2, or with the wild-type DH5α strain. B) Percentage of events in Q2 for combinations of yeast displaying IB10, Nb2, or the nondisplaying wild-type RJY100 strain with *E. coli* displaying intimin, Ag2, or the nondisplaying wild-type DH5α strain. Error bars represent the standard deviation of 3 biological replicates for yeast displaying nanobodies.

### Model system supports evaluations of parameters that affect cell-cell interactions

To differentiate between specific and nonspecific cell-cell interactions, coincubation conditions such as total cell density, yeast to bacteria cell ratio, and incubation time may need to be varied depending on parameters including the affinity of the binding molecule to the target molecule and surface display densities on both yeast and bacterial cell surfaces. In the experiments described above, the number of displayed constructs on the yeast surface using the galactose-inducible Aga1p-Aga2p fusion system is approximately 3 × 10^4^ fusion molecules per cell.^*24*^ The number of antigens on the *E. coli* surface displayed using the scheme employed in this work has not been quantified to our knowledge, but a rough estimate is 3.6 × 10^4^ molecules per cell for aTc induction of intimin-anchored proteins.^*25*^ The K_d_ of the Nb2-Ag2 interaction, which led to reliable cell-cell interactions in comparison to background levels, has been previously reported to be 190 ± 30 nM at room temperature.^*26*^ Importantly, as shown in the dot plots for representative populations of yeast displaying nanobodies labeled for N- and C-terminal detection, the expression of full-length nanobodies is not homogenous. Consistent with prior studies utilizing the galactose-inducible Aga1p-Aga2p display system, some cells appear to possess higher surface-densities of nanobodies available for interaction (Figure S2).^*18, 27*^ These observations motivate further exploration of parameters that can enable discrimination between cell-cell interactions mediated by specific binding and cell-cell interactions mediated by nonspecific binding.

As a step towards investigating a wider range of coincubation conditions, we attempted to modulate the surface density of nanobodies present on the yeast surface to better understand the role of ligand surface density (or avidity) in this system. Previous work has shown that ligand density can greatly influence target binding and enrichment.^*28, 29*^ To control ligand display levels, we treated yeast cells with varying concentrations of dithiothreitol (DTT), which can have the combined effects of 1) reducing the two disulfide bridges linking Aga1p and Aga2p to dissociate the displayed protein from the cell surface^*11*^ and 2) reducing the disulfide bridges within the displayed protein to change it to a conformationally inactive state.^*30*^ In independently conducted experiments of yeast and *E. coli* surface display of three interaction pairs—IB10-intimin, Nb2-Ag2, and Nb3-Ag3—we determined that the nanobody display level (and therefore surface density) was linearly correlated (p-values = 0.0009, 0.0047, and 0.0007, respectively) with the frequency of detected interaction events; higher display levels led to higher percentages of interactions (Figure S5).

The use of longer linkers may also change yeast-bacteria interaction strengths by altering the accessibility of displayed ligands. The linker length used in the above described experiments is 40 amino acids. A recent study using a yeast-mammalian cell interaction system investigated the use of an intermediate linker (80 amino acids) and long linker (641 amino acids) for cell-cell interaction experiments. The study found that the long linker allowed for higher enrichment, likely due to improved surface accessibility, although the surface density of displayed ligands and percent of ligand-displaying cells were lower for cells displaying the long linker compared to the intermediate linker.^*10*^ We investigated cell-cell interactions with the IB10-intimin and Nb2-Ag2 nanobody-antigen interaction pairs using both the 80- and 641-amino acid linkers. In our hands, we observed an increase in cell-cell interactions for the cognate pairs as well as an increase in nonspecific cell-cell interactions with the longer linkers. Based on these data, it was not possible to determine if the longer linkers significantly enhanced nanobody-antigen interactions under these conditions (Figure S6). Further exploration of the effects of these parameters on cell-cell interactions are warranted in a future study to fully understand the advantages and limitations of the yeast-bacteria interaction assay.

## Conclusions

In this study, we demonstrate that flow cytometry can be used to identify yeast-bacteria interactions mediated by binding of surface available cognate nanobody-antigen pairs. Furthermore, we validate the use of fluorescent dyes to stain cells used in this assay and show representative microscopy images of yeast-*E. coli* cell-cell complexes. These cell-cell complexes are stable enough to be enriched by FACS, as determined by both flow cytometry and sequencing of sorted populations. Yeast surface display densities and linker lengths were shown to play important roles in cell-cell interactions mediated by specific binding events. The effects of these parameters on yeast-*E. coli* cell-cell interactions for screening applications warrants further assessment in future studies. Finally, we test a biologically relevant application of this interaction system towards a nanobody-antigen interaction in a model of EHEC adhesin (i.e., intimin) overexpression and show that the cell-cell interaction assay is able to discriminate between the cognate IB10-intimin interaction and non-specific interactions.

Based on these results, this yeast-bacteria interaction system could potentially be used to discover surface-binding ligands towards any bacterial species amenable to laboratory cultivation, fluorescence staining, and flow cytometry. A yeast display system offers many advantages for ligand discovery in flow cytometry format,^*20, 31*^ including: 1) a eukaryotic translation apparatus that facilitates biosynthesis of complex polypeptides including genetic encoding of different chemical functionalities (e.g. natural post-translational modifications such as disulfide bond formation and glycosylation, noncanonical amino acid (ncAA) substitutions for bioorthogonal chemistries and crosslinking^*32*^), and other chemical groups amenable to selective modification on the yeast surface;^*33, 34*^ 2) a robust cell wall to facilitate straightforward handling; 3) efficient propagation relative to other eukaryotic organisms; and 4) lack of susceptibility to many antibacterial compounds.^*35*^ Bacterial surface-binding molecules have broad applications for biofilm control,^*4*^ therapeutic inhibition of bacterial adhesion to host cells,^*36*^ and selective targeting of bacterial species within a microbial community.^*37*^ The yeast display-based cell-cell interaction assay described in the present study provides a flexible platform to discover and characterize these binding molecules.

## Materials and Methods

### Materials

All DNA fragments and oligonucleotides for molecular cloning and sequencing were purchased from Genewiz. Restriction enzymes were purchased from New England Biolabs. Epoch Life Science miniprep kits were used for *E. coli* plasmid purification. Materials used for yeast plasmid purification are described in the methods section *Preparation for amplicon sequencing and data analysis*. Calcofluor white was purchased from Sigma-Aldrich and SYTO 9 was purchased from Thermo Fisher Scientific.

### Plasmid construction

Previously characterized orthogonal nanobody-antigen interaction pairs (Nb1-Ag1, Nb2-Ag2, Nb3-Ag3) used in an *E. coli*-*E. coli* interaction format^*15*^ were adapted to a yeast-*E. coli* interaction format by cloning the nanobodies into a pCTcon2 yeast display vector. The set of previously designed and characterized nanobody-antigen interaction pairs adapted to this study are Nb1 (anti-Akt3PH)-Ag1 (Akt3PH), Nb2 (anti-EPEA)-Ag2 (EPEA), and Nb3 (anti-P53TA)-Ag3 (P53TA).^*15*^ Plasmids for *E. coli* surface display of Nb1, Ag1, Nb2, Ag2, Nb3, and Ag3 were obtained from Addgene: pDSG372 (Addgene plasmid #115602), pDSG358 (Addgene plasmid #115599), pDSG375 (Addgene plasmid #115603), pDSG419 (Addgene plasmid #115600), pDSG398 (Addgene plasmid #115604), pDSG360 (Addgene plasmid #115601). The pCTcon2-nanobody constructs were cloned by PCR amplification of the nanobody genes with the oligonucleotides listed in Table S1, digestion of the pCTcon2 vector with NheI and BamHI (New England Biolabs), and Gibson assembly. The IB10 nanobody^*6*^ was synthesized as a double-stranded, linear DNA fragment (FragmentGENE, Genewiz) and also cloned by Gibson assembly into pCTcon2. pDSG_Intimin_full_length was provided by the Mougous lab at the University of Washington.^*37*^ pCT80 and pCT641 were provided by the Hackel lab at the University of Minnesota.^*10*^

### Culture conditions

RJY100 yeast cells were transformed with the pCTcon2 nanobody-display vectors. Yeast containing the pCTcon2 nanobody-display vector were grown in 2 ml SD-SCAA −Trp −Ura +pen/strep pH 4.5 in 15 ml polypropylene tubes at 30°C in a shaking incubator overnight to saturation, diluted to OD = 1 in 2 ml of the same medium, grown for an additional 2–3 h at 30°C to approximately OD = 2. Pen/strep was used at a 1× total concentration from a 100× stock solution, containing 10000 IU penicillin and 10000 µg/ml streptomycin (Corning). An aliquot of each culture corresponding to an OD of approximately 1 in a 4 ml volume was spun down. The supernatant was aspirated, and the cell pellet was resuspended in 4 ml SG-SCAA −Trp −Ura +pen/strep. The cultures were induced by incubation in a 20 °C shaking incubator for 16 hours. Three biological replicates were performed for each of Nb1, Nb2, and Nb3-displaying cells by picking three different colonies on the yeast transformation plates and treating them the same throughout the experiment. Wild-type RJY100 was grown in SDSCAA −Ura +pen/strep or SGSCAA −Ura +pen/strep following the corresponding steps above. Full-length surface display was assessed by labeling both the N-terminal hemagglutinin (HA) and C-terminal c-Myc tag flanking the displayed nanobody as described in a previous study.^*38*^

One *E. coli* colony transformed with each of Ag1, Ag2, Ag3 were grown in 2 ml LB kan (50 µg/ml kanamycin) at 37 °C in a shaking incubator overnight to saturation, diluted to OD 0.05 in 4 ml LB kan, grown for an additional 2–3 hours to an OD of approximately 0.6, then induced with a final anhydrotetracycline (aTc) concentration of 200 ng/ml. The cultures were induced for approximately 16 hours. Wild-type DH5α was grown in LB, otherwise following the same steps above. ATc was prepared by making a 1 mg/ml stock solution in ethanol, then diluting it 1:10 in water to make a 100 µg/ml working stock solution.

### Fluorescence staining

After induction, the OD of each culture was measured using a cuvette with a 1 cm path length by diluting the culture 1:10 in PBS pH 7.4 (Note: Precise OD measurements are critical for these experiments since the frequency of nonspecific interactions are correlated to the ratios of yeast: *E. coli*. See Figure S1 for details). A corresponding volume of each culture required to prepare a 2 ml volume at OD 1 was pipetted from each tube and pelleted. Each cell pellet was resuspended in 2 ml PBS. All centrifugation steps were performed at 5000 rpm (corresponding to approximately 2000 × g) for 2 minutes on an Eppendorf 5424 tabletop centrifuge, unless noted otherwise. Calculations assume 1 × 10^9^ cells/ml *E. coli* at OD 1 and 3 × 10^7^ cells/ml yeast at OD 1.^*39*^

To stain yeast, 2 µl of a 1 g/L stock solution of calcofluor white (CFW) was added to each of the 2 ml volumes (1 µl per OD1/ml) and incubated for at least 15 minutes. To stain *E. coli*, 2 µl of a 3.34 mM stock solution of SYTO 9 in DMSO (LIVE/DEAD® BacLight™ Bacterial Viability and Counting Kit) was added to each of the 2 ml volumes (corresponding to 1 µl SYTO9 per 1 ml of cell suspension at OD 1) and incubated in the dark for at least 15 minutes. For microscopy, 4× higher cell densities were used for practical reasons: the density at which cell complexes were optimally detected above background on the flow cytometer is not suitable for microscopy because the cells are too sparsely distributed on the agarose pads. The tubes were briefly mixed, then incubated on a rotary wheel for 15 minutes while covered from light. *E. coli* cells were washed by pelleting the cells, aspirating the supernatant, and resuspending in PBS, twice. SYTO 9 was previously shown to stain yeast cells, whereas CFW did not significantly stain *E. coli* at the concentrations used, so no wash step was deemed necessary for CFW-stained yeast prior to coincubation with *E. coli*.

### Flow cytometry

Flow cytometry analysis was performed on an Attune NxT flow cytometer. Relevant flow cytometer settings were: 25 µl/min flow rate, 10000 events collected per sample, FSC:1, SSC:200, BL1, RL1, VL1: 250. The minimum FSC voltage was used since previous experiments determined that higher FSC voltages caused many events to appear outside the axis limits for coincubated cultures, especially for those with an expected interaction. Though the FSC scatter data captures the same trends as reported in fluorescence data since a higher FSC can indicate formation of large cell aggregates (data not shown), these data exhibit high coefficients of variation (CVs) associated with low voltage settings. Therefore, FSC scatter data was not used in analyses reported here. Data analysis was performed using FlowJo and Microsoft Excel.

### Microscopy

Microscopy was performed using a DMi8 automated inverted microscope (Leica Microsystems, #11889113) equipped with a CCD camera (Leica Microsystems, #DFC300 G), and LED405 (Ex 375-435 nm, Em 450-490 nm, exposure time – 30 ms, gain 3.3) and YFP (Ex 490–510 nm, Em 520–550 nm, exposure time – 30 ms, gain 3.3) filter cubes. 1 µl of coincubation cultures were spotted on agarose pads (2 % w/v, 1 mm thick).^*40*^

### Fluorescence-activated cell sorting

Cell sorting was performed on a Bio-Rad S3e Cell Sorter. 1:1:1, 10:1:10, and 100:1:100 mixtures of Nb1:Nb2:Nb3 displaying yeast cells were prepared by normalizing ODs and adding corresponding volumes to achieve the indicated ratios. Sorting gates were drawn on dot plots with FL1 and FL4 on the axes, with gates positioned to capture events exhibiting high levels of fluorescence in the FL1 green emission channel. The FL4 far-red emission channel was chosen since the S3e sorter model used in this work is not equipped with a laser capable of exciting CFW. FL4 has minimal overlap with the emission spectrum of SYTO 9. The same general procedure as in the analytical flow cytometry experiment was followed for quantification of cell-cell interactions. Pre-sort populations (prior to mixing with *E. coli*) were saved for yeast miniprep and amplicon sequencing. After sorting, cells were recovered in SD-SCAA ™Trp ™Ura +pen/strep pH 4.5 for 2 days. Post-sort populations were saved and coincubation assays were performed concurrently with single and mixed ratios of nanobody-displaying yeast as controls.

### Preparation for amplicon sequencing and data analysis

Yeast miniprep was performed using the E.Z.N.A.® Plasmid DNA Mini Kit I (OMEGA Bio-tek, #D6943-02) with slight modifications. Approximately 5-10 OD*ml cells were resuspended in solution I (250 µl) supplemented with lyticase (1000 U, Sigma, catalog# SAE0098-20KU) and incubated at 37 °C for 1 h. The remaining steps were performed according to the manufacturer’s protocol. The extracted DNA was quantified using a spectrophotometer and approximately 100 ng was used as template for PCRs. All samples were prepared for amplicon sequencing using unique, barcoded primers flanked by Illumina sequencing adaptors (Table S1). 15–20 cycles of PCR were performed to amplify the region spanning the nanobody using the primers listed in Table 1. The barcoded samples from triplicate experiments were pooled and sent for amplicon sequencing (2×250 bp, Genewiz, New Jersey, USA). For each pooled sample, we received approximately 100,000 sequencing reads. These data were processed according to a previously described bioinformatic workflow using Geneious Prime® 2020.2.4.^*41*^ Briefly, the reads were paired and merged using the BBMerge package and filtered for poor-quality reads using the BBDuk package. The reads were then mapped onto the reference genes (Nb1, Nb2, and Nb3) using BowTie2.^*42*^ The number of reads mapped to Nb1, Nb2 and Nb3 were used to estimate the fold enrichment.

## Supporting Information

Figures S1-S6 show the effect of cell ratios on detected interaction events, quantification of full-length nanobody display, sorting gates used for FACS, calibration of detected interaction events to the proportion of cognate nanobody-displaying cells within a population, effect of DTT modulation of surface display density and total displayed nanobodies, and effect of linker length. Table S1 lists all DNA sequences relevant to the methods in this study.

## Acknowledgments

This work was supported in part by a grant from the National Institute of General Medical Sciences of the National Institutes of Health (1R35GM133471 to J.A.V.) and the Karol Family Professorship (to K.L.). The content is solely the responsibility of the authors and does not necessarily represent the official views of the National Institutes of Health or Tufts University. Plasmids pCT80 and pCT641 were a gift from the laboratory of Benjamin Hackel. Plasmid pDSG_Intimin_full_length was a gift from the laboratory of Joseph Mougous.

