## Supplementary Information for "Flow Cytometric Evaluation of Yeast-Bacterial Cell-Cell Interactions"

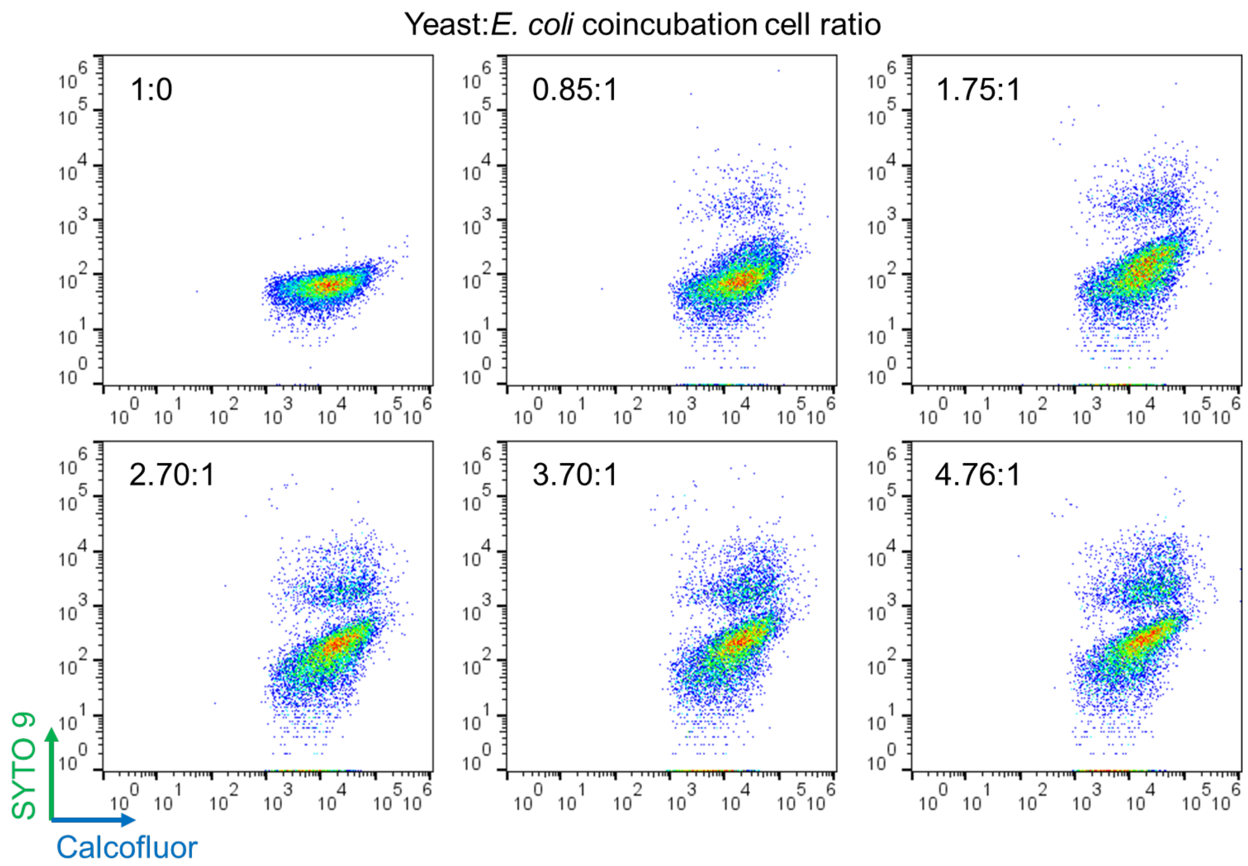


**Figure S1.** Different coincubation ratios of wild-type yeast (RJY100) to *E. coli* (DH5α) cells measured on the flow cytometer.

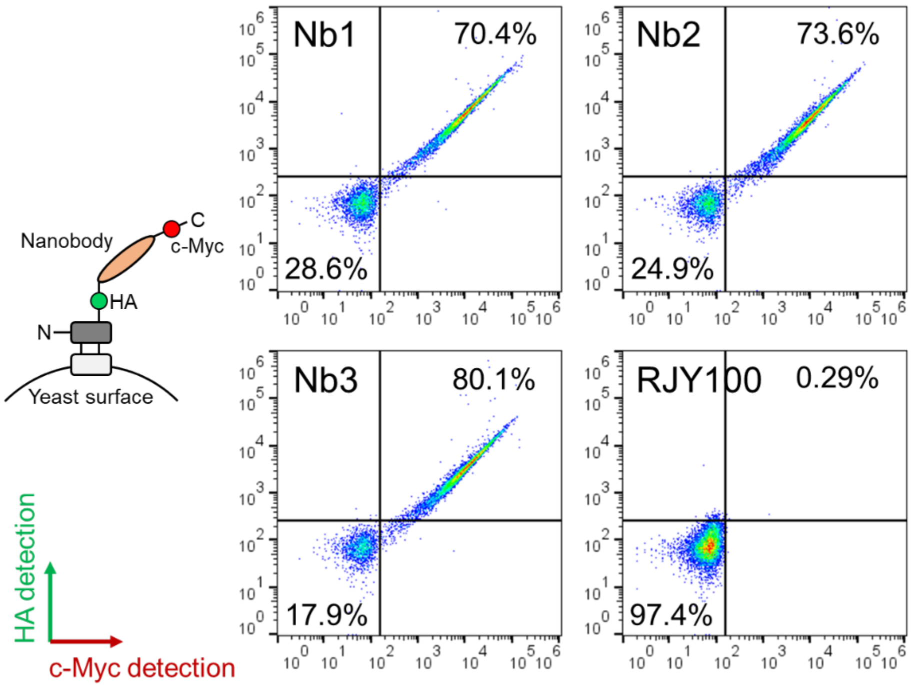


**Figure S2.** Representative dot plots for N- and C-terminal labeling of HA and c-Myc tags, respectively, in nanobody-expressing yeast clones, and the wild-type RJY100 with no surface display.

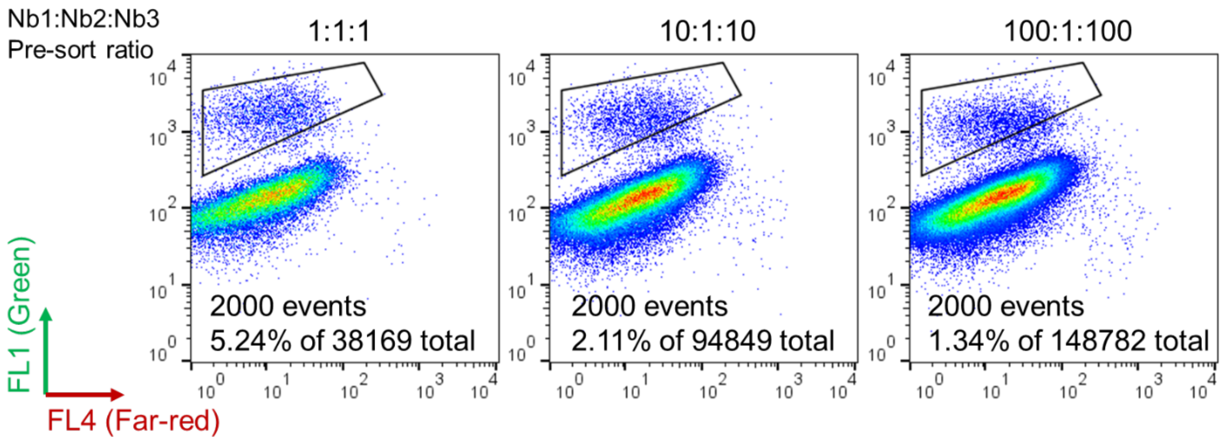


**Figure S3.** Representative dot plots of sorting gates containing 2,000 sort counts each, acquired using an S3e cell sorter.

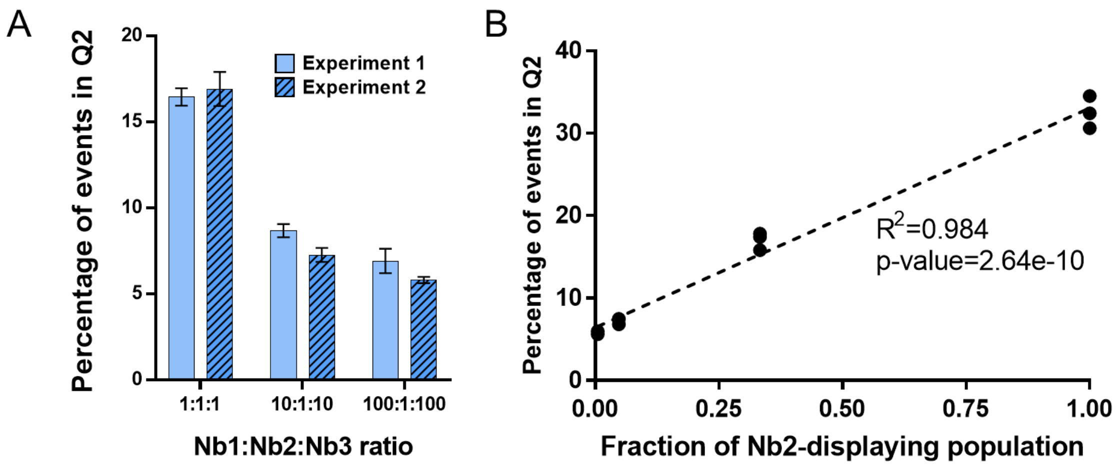


**Figure S4.** Percentage of events in Q2 (high blue and green fluorescence) for mixed nanobody ratios of Nb1, Nb2, and Nb3 (1:1:1, 10:1:10, 100:1:100) coincubated with Ag2-displaying *E. coli*. A) Experiments run on separate days as pre-sort and post-sort controls. B) Calibration curve for fraction of Nb2-displaying population versus percentage of events in Q2.

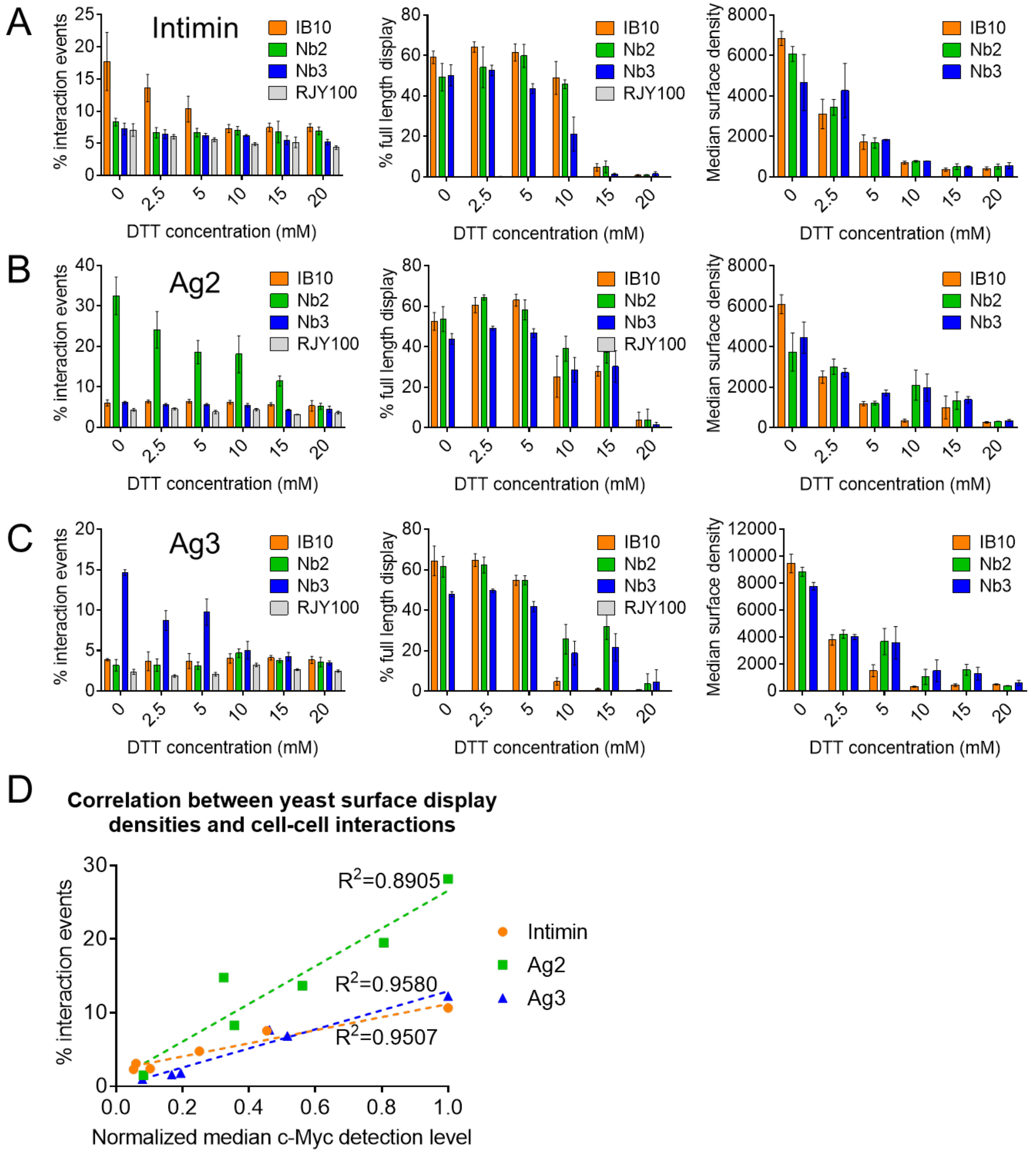


**Figure S5.** Effect of DTT treatment on surface display mediated cell-cell interactions between yeast and *E. coli* cells. The samples for each set (coincubation with *E. coli* displaying intimin, Ag2, or Ag3) were processed at 4-hour intervals and each panel can be considered the result of independent experiments. The fluorescence values of different panels cannot be directly compared due to the long experimental processing time required to assay biological triplicates. During this time, changes in cell physiology and DTT degradation can possibly become confounding factors. Panels

A, B, and C each contain plots for: 1) Measured cell-cell interaction events quantified as the percentage of events that exhibit high levels of both blue and green fluorescence. 2) Amount of displayed full-length nanobodies quantified as the percentage of events with both green and red fluorescence after labeling of the N- and C- terminal HA and c-Myc tags, respectively. 3) Surface display density quantified as the median red fluorescence intensity of the full-length nanobody displaying population. A) Coincubation of different yeast transformants with *E. coli* displaying intimin. B) Coincubation of different yeast transformants with *E. coli* displaying Ag2. C) Coincubation of different yeast transformants with *E. coli* displaying Ag3. D) Correlation between normalized median red fluorescence intensity of full-length nanobody displaying cells and the baseline subtracted percentage of interaction events for all three experiments.

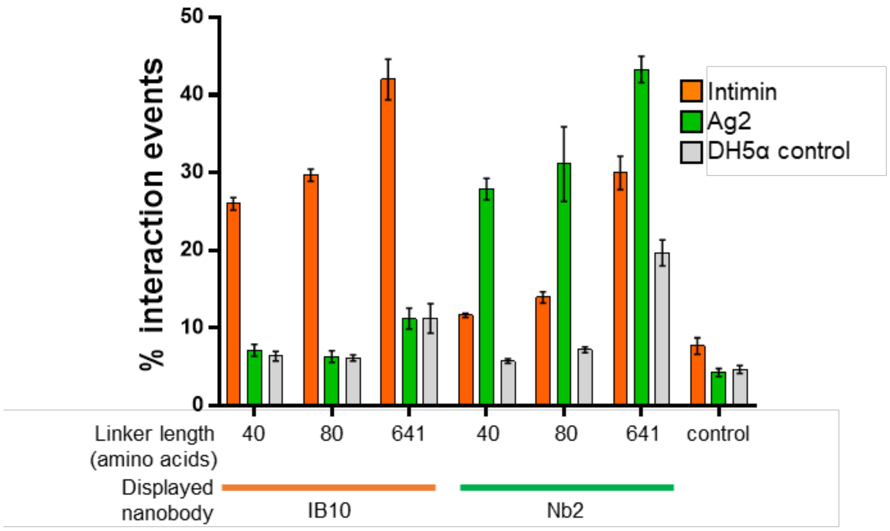


**Figure S6.** Percentage of interaction events detected for different nanobody-antigen interaction pairs with varying linker lengths used for nanobody display on the yeast surface.

**Table S1.** List of DNA sequences used for cloning and amplicon sequencing. For amplicon sequencing primers: Red = sequencing adaptor; Purple = barcode; Black lowercase = target binding region.

| Cloning | |
|---|---|
| Nb1-Fwd | GTGGAGGCGGTAGCGGAGGCGGAGGGTCGGCTAGCCAAGTTCAGTTGCAAGAGAGTGGAG |
| Nb1-Rev | CCTCTTCAGAAATAAGCTTTTGTTCGGATCCACTACTAACGGTAACTTGCGTTCCC |
| Nb2-Fwd | GTGGAGGCGGTAGCGGAGGCGGAGGGTCGGCTAGCCAAGGCCAGCTTGTGGAATC |
| Nb2-Rev | CCTCTTCAGAAATAAGCTTTTGTTCGGATCCTGAGGATACCGTCACCTGTGTG |
| Nb3-Fwd | GTGGAGGCGGTAGCGGAGGCGGAGGGTCGGCTAGCCAAGTTCAGTTACAGGAATCTGGG |
| Nb3-Rev | CCTCTTCAGAAATAAGCTTTTGTTCGGATCCGCTACTCACCGTAACCTGTGTC |
| IB10 FragmentGENE | GTGGAGGCGGTAGCGGAGGCGGAGGGTCGGCTAGCATGGCTCAAGTTCAATTGGTTGAATC<br>TGGTGGTGGTTCTGTTCAAGCTGGTGGTTCTTTGAGATTGTCTTGTAAAGCTTCTGGTTTTACT<br>TTTCCATATTCTGATATGGGTTGGTATAGACAAACTCCAGGTAATGAATGTGAATTGGTTTCT<br>ACTACTGGTCCAGAATCTGATCCATCTTTTTGGTATGCTGATTCTGTTAAAGGTAGATTTACTA<br>TTTCTAGAGATAATACTAAAAATACTGTTTATTTGCAAATGAATGATTTGAAACCAGAAGATA<br>CTGGTATGTATGTTTGTGCTTCTGAATTGGGTGCTGGTTCTGGTAGATGTTATGGTTATCATT<br>ATTGGGGTCAAGGTACTCAAGTTACTGTTTCTTCTGCTGCTGGATCCGAACAAAAGCTTATTT<br>CTGAAGAGGACTTGTAATAGCTCGAGATCTGATAACAACAGTGTAGATGTAACAAA |
| Amplicon sequencing | |
| Forward primer-1 | ACACTCTTTCCCTACACGACGCTCTTCCGATCTAGCGATTA**ggcggagggtcggctagcc** |
| Forward primer-2 | ACACTCTTTCCCTACACGACGCTCTTCCGATCTAGGAATTG**ggcggagggtcggctagcc** |
| Forward primer-3 | ACACTCTTTCCCTACACGACGCTCTTCCGATCTGGCTCAAC**ggcggagggtcggctagcc** |
| Forward primer-4 | ACACTCTTTCCCTACACGACGCTCTTCCGATCTGTTGATGG**ggcggagggtcggctagcc** |
| Forward primer-5 | ACACTCTTTCCCTACACGACGCTCTTCCGATCTAGTTATAC**ggcggagggtcggctagcc** |
| Forward primer-6 | ACACTCTTTCCCTACACGACGCTCTTCCGATCTACATTCCA**ggcggagggtcggctagcc** |
| Reverse primer | GACTGGAGTTCAGACGTGTGCTCTTCCGATCT**ttcagaaataagcttttgttcggatcc** |
